# Deciphering the Metabolic Shifts in The Hippocampus of Mice Subjected to Near Low Dose Radiation: Insights from Metabolomics and Integrated Multi-omics

**DOI:** 10.1101/2023.11.05.564910

**Authors:** Rekha Koravadi Narasimhamurthy, Babu Santhi Venkidesh, M B Joshi, Bola Sadashiva Satish Rao, Krishna Sharan, Kamalesh Dattaram Mumbrekar

## Abstract

Recent years have witnessed a drastic upsurge in neurological disorders, with sporadic cases contributing more than ever to their cause. Radiation exposure through diagnostic or therapeutic routes often results in neurological injuries indicative of neurodegenerative pathogenesis. Nevertheless, the impact of low doses of radiation on the brain remains a subject of extensive discussion, as research findings have presented conflicting evidence regarding potential harm and benefits. In the present study, C57/BL mice were exposed to a whole-body single dose of 0.5 Gy X-ray. Fourteen days after treatment, the animals were euthanized, and the hippocampus was isolated and processed for metabolomic analysis. Statistical and bioinformatic analysis revealed 115 metabolites altered in the radiation-exposed group, while pathway enrichment analysis unveiled alterations in tyrosine, phenylalanine, aminoacyl-tRNA metabolism, arginine biosynthesis, glutathione, arginine, proline metabolism, etc. Furthermore, a multiomics interaction network of the genes and the metabolites was constructed to gather an overview of their interaction with the neighboring genes and metabolites in different pathways. These metabolic pathways correlate with synthesizing neurotransmitters such as dopamine and neurodegenerative diseases such as Alzheimer’s, Parkinson’s, and dementia. The present study findings unveiled metabolomic level regulation of low-dose radiation-induced neurotoxicity and its implication in the pathogenesis of neurological disorders.

## Introduction

Nervous system disorders are one of the leading causes of suffering and death worldwide and contribute vastly to their rising global burden. Owing to their multifactorial origin, their various causative mechanism is largely unexplored, except in cases where the basis is familial. Studies have shown that environmental exposure to neurotoxicants can alter several metabolite levels in the brain and induce neurodegeneration (Rodrigues et al. 2022). Radiation exposures have been linked to the triggering of sporadic cases of neurodegeneration through the accumulation of chronic damage over the early years and manifesting as disease during old age (Sharma et al. 2018). In the brain, the hippocampus is the site of neurogenesis and controls various complex behavioral capabilities, including cognition, memory, and learning, exploratory behavior, and emotions (Peng and Bonaguidi 2018). Further, it is a vulnerable and plastic region that is sensitive to various stimuli and dynamically alters its morphology and cellular and molecular functions (Anand and Dhikav, 2012). The hippocampus is a well-known target for neurotoxic insults, particularly in causing metabolomic perturbations (Walsh and Emerich 1988). Moreover, in both post-natal rodents and adults, the subgranular and the subventricular zones of the hippocampus harbor neural progenitor cells that are known to be radiosensitive, employed with the aim of replenishing damaged or dead neural cell population (Katsura et al. 2016).

Ionizing radiation (IR) is commonly employed in diagnosing and treating various diseases. Over the years, the number of individuals undergoing procedures involving IR has increased, leading to increased studies investigating its long-term health effects (Smith-Bindman et al. 2008; Lin 2010). Furthermore, in patients undergoing radiotherapy for brain-related cancers, it has been shown that some portion of surrounding normal tissue can also inadvertently be exposed to lower doses of IR, making it a cause of concern (Auerbach et al. 2023). Such procedures spanning multiple exposures can lead to accumulated damage that can manifest as cognitive changes, neurological sequelae, or even cancer in some cases (Collett et al. 2020; Narasimhamurthy et al. 2022). High-resolution metabolomic profiling has been a recommended technique for identifying potential biomarkers involved in radiation response.

It facilitates better translation of radiation-associated pathogenesis post-exposure (Menon et al. 2016). Previously, studies have been carried out to understand the metabolomic changes in glioblastoma patients undergoing radiotherapy (Wibom et al. 2010) or at higher doses (10Gy and 30 Gy) in mouse models using serum/plasma samples (Meng et al. 2022) or the hippocampus (Torres et al. 2019). One of the early studies involving a metabolomics approach following cranial irradiation of 8 Gy revealed several alterations in metabolites in the citric acid cycle, neurotransmitter metabolism, and glutamate metabolism in the hippocampus (Rana et al. 2014). Mice irradiated with whole body 0.5 Gy of ^1^H irradiation or ^16^O irradiation exhibited alterations in several pathways such as amino acid metabolism, tyrosine metabolism, lysine metabolism, glycolysis, glycerophospholipid metabolism, gluconeogenesis, etc. in plasma samples (Dissmore et al. 2021). Another study investigating the changes in serum profiles after whole brain irradiation (5 × 2 Gy) reported alterations in the branched-chain amino acids valine, isoleucine, and leucine (Maddens et al. 2020). Pazzaglia et al. observed transient metabolomic changes in mice at 0.1 Gy, while 2 Gy X-ray induced changes in the TGF-β and DAG/IP3 pathways, while also affecting synaptic plasticity and other neuronal functions (Pazzaglia et al. 2021). Wistar rats irradiated with 0.14 Gy of carbon (^12^C) nuclei showed suppressed dopamine turnover in several regions of the brain except in the hippocampus, where the contrast was observed, with decreased levels of norepinephrine in the amygdala (Kokhan et al. 2022). It is important to note that a complete assessment of the entire spectrum of changes cannot be achieved using only a single omic approach, making it essential to employ multiple platforms to identify the most comprehensive set of potential biomarkers and their interactions (Gonzalez-Riano et al. 2016). In view of the increasing low dose exposure during diagnostic exposures (Brenner 2014) and space exploration (Roy-O’Reilly et al. 2021), it is important to study the changes in brain metabolome and its relevance with neurodegenerative disease pathways. Further, the integrated analysis of transcriptomic and metabolomic data can aid in understanding the multiomics regulation of neurotoxic effects and predict how they can eventually pose a risk in the induction of neurodegeneration. In addition, understanding the metabolites and the altered pathways will give better insight into understanding response to low-dose IR.

Our study is focused on understanding the acute metabolomic response to radiation in younger aged mice (4-5 weeks) in the hippocampus at near-low dose exposure to 0.5 Gy, which has not been done before. Understanding the metabolomic alterations of the hippocampus, in particular, is crucial in characterizing their potential involvement in other molecular and functional aspects. Thus, the present study illustrates the alterations in the metabolomic landscape of the hippocampus post-individual exposure to 0.5 Gy IR. We then correlate these changes to potentially altered pathways that are critical in neuronal functioning and disease regulation. Furthermore, integration of the metabolomic data with previously obtained transcriptomic data (Narasimhamurthy et al. 2023) in the same set of animals is also carried out to understand how multi-omics analysis can shed enhanced insight into its regulatory mechanisms.

## Materials and Methods

### Animal maintenance and treatment

C57BL/6 mice (approximately 5 weeks old and weighing approximately 27 g) were maintained in the Central Animal Research Facility, MAHE, Manipal, with prior ethical approval from the Institutional Animals Ethics Committee, MAHE, Manipal (IAEC/KMC/108/2019). Animals were maintained in a controlled environment with a temperature of 20°C ± 2°C and a 12 h light and dark cycle. Nine animals in two groups each, control and radiation (0.5 Gy) were taken for the study wherein the animals in the radiation group were subjected to 0.5 Gy whole-body single-dose radiation with the Versa-HD Linear Accelerator (Elekta, Sweden) using 6 MV Photons while the control animals were sham irradiated. Following nine days of treatment and behavioral assays for four days, the animals were euthanized by cervical dislocation, and the hippocampus was isolated and stored at -80°C until further processing.

### Sample preparation for Mass spectrometry

Hippocampus tissues were weighed and accordingly minced in methanol and water (4:1, v/v) at a final concentration of 20 mg/ml. Samples were then briefly vortexed for 30 s, snap-frozen using liquid nitrogen, and then sonicated for 5 min in 3 cycles. The samples were incubated at -20°C for 1 hour, followed by centrifugation at 12000 rpm for 10 min. The supernatant was then collected and lyophilized, and the residue was redissolved in 100 μL of cold acetonitrile and water (50:50, v/v) containing 1% formic acid. The samples were again centrifuged at 12,000 rpm for 10 min, and the supernatant was collected and stored at -80°C until LC/MS injection.

### LC/MS conditions

Mass spectrometry was carried out using ESI-QTOF (Agilent 6250 TOF-MS, Agilent Technologies, USA) coupled with a high-performance liquid chromatography system (Agilent 1200 series, USA). The sample was reconstituted in 100μL of water: acetonitrile containing 0.1% formic acid. The analysis of the samples was performed in positive mode with an injection of 5 μL of sample per run into the Agilent analytical column (ZORBAX Eclipse XDB C18, 4.4×250 mm, 5 microns). The mobile phase used was acetonitrile and water with 0.1% formic acid. The conditions of the mass spectrometry run were as follows: a gas temperature of 250°C, gas flow of 8 L/min, nebulizer pressure of 40 psig, and ESI capillary voltage of 3500 V. Samples were run in triplicate with each run lasting for 45 minutes.

### Metabolomic data analysis

The raw data obtained in .d format were converted to mzML format using the tool MS Convert. The mzML converted files were then uploaded to Metaboanalyst 5.0 (Pang et al. 2021), where the sample data were checked for their integrity, and spectral processing and annotation were carried out in the positive mode using a tolerance limit of ±15 ppm. After data filtering and normalization, log transformation and autoscaling were performed. Fold change analysis and t-tests were carried out to identify the list of downregulated and upregulated metabolites. Partial Least Squares Discriminant Analysis (PLS-DA), a statistical tool that determines and classifies the metabolites that contribute the most to the separation between the two groups, allowing for the recognition of potential biomarkers, was also carried out. This was done by attributing a VIP metric score to the metabolites based on the metabolite measurements. Further, PLS-DA was also performed to identify statistically distinct metabolomic features, and their association with various pathways and disease conditions was assessed using pathway enrichment analysis.

### Metabolite-gene interaction analysis

The HMDB IDs of the metabolites were input into the online tool Metabridge (Blimkie et al., 2020), and a list of genes regulating the metabolite/enzyme levels was obtained. The list of genes was compared with our published hippocampal transcriptomic data (Narasimhamurthy et al. 2023) to find commonly altered genes. Additionally, we also used Metscape (Karnovsky et al. 2012), a plugin in Cytoscape (Shannon et al. 2003), to find a direct correlation between the differentially expressed genes from our experiment with the list of metabolites identified and the pathways that they are involved in.

### Statistical analysis

**A** t-test was carried out to individually compare radiation-exposed treatment groups with respect to control. A p-value < 0.05 was considered to be significant.

## Results

Comparison between the control and radiation groups revealed 71 upregulated and 52 downregulated metabolites in the radiation group with a fold enrichment filter of 2 (Fig 1a and table 1). Principal component analysis identified metabolomic features displayed maximum variation between the two groups, with components one and two displaying 70.8% and 16.9% variation, respectively (Fig 1b).

**Fig 1:**
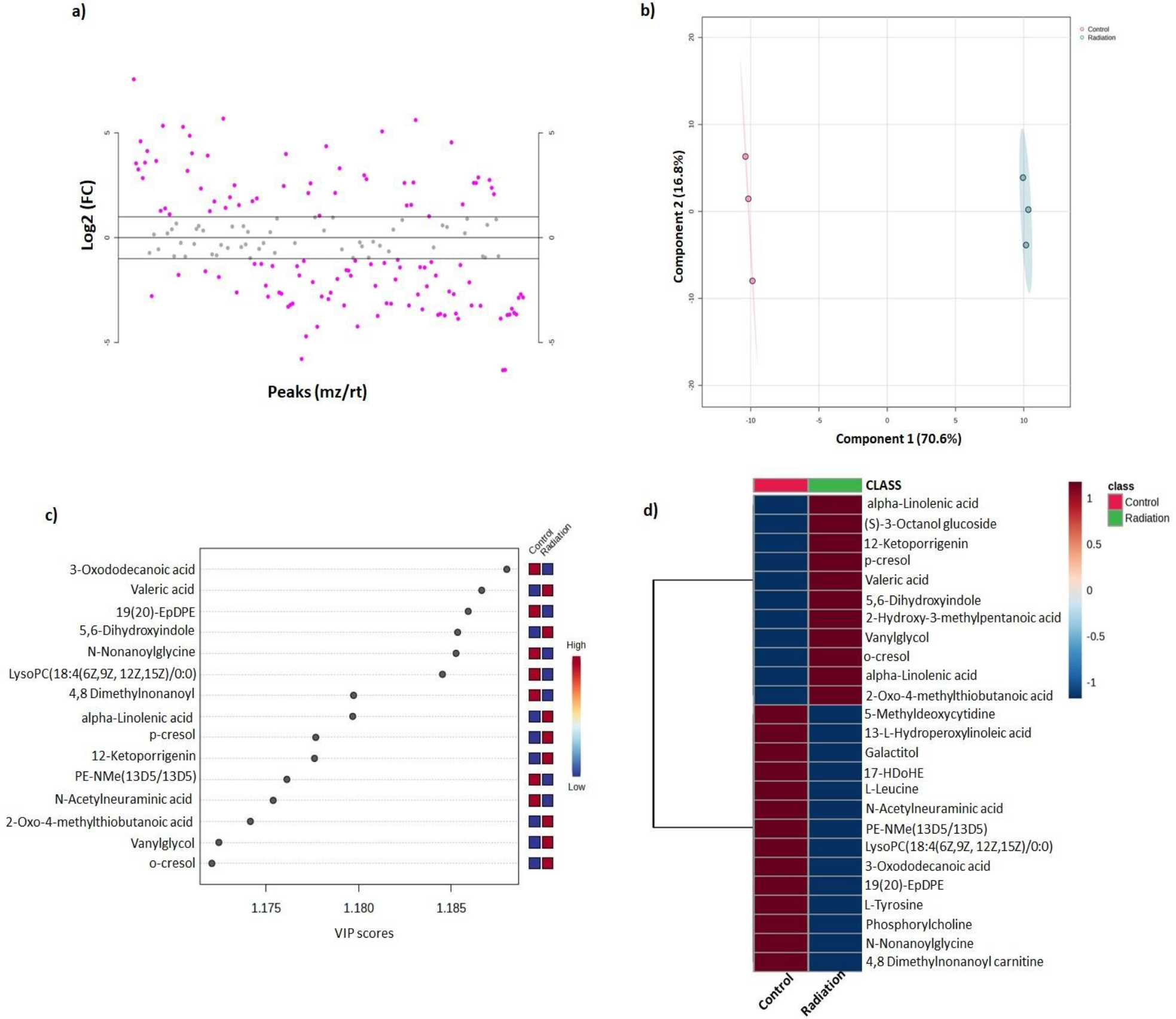
Effect of low-dose radiation exposure on hippocampal metabolites. a) Number of upregulated and downregulated metabolites; b) Principal component analysis showing the variation in the data; c) PLS-DA analysis depicting the top discriminating metabolites; d) Top 25 metabolites and their levels post low-dose radiation exposure

### PLS-DA analysis

The PLS-DA analysis showed that the 3-oxododecanoic acid, valeric acid, 19,20-epoxydocosapentaenoic acid, 5,6-dihydroxyindole, and N-nonanoylglycine were the top 5 discriminating features between the control and radiation groups according to their VIP scores (Fig 1c). Cluster heatmap depicted valeric acid, 5,6-dihydroxyindole, (S)-3-octanol glucoside, 2-oxo-4-methylthiobutanoic acid, and 2-hydroxy-3-methylpentanoic acid were the top five most downregulated metabolites while, lysoPC(18:4(6Z,9Z,12Z,15Z)/0:0), lysoPA(24:1(15Z)/0:0), N-acetylglutamine, N-acetyl-L-glutamic acid, and 5Z-dodecenoic acid, were the top five most upregulated metabolites in radiation (Fig 1d). Furthermore, the normalized fold change values of some of the critical metabolites involved in neurodegenerative disease pathways, compared to the control, were represented in box whisker plots (Fig 2a and b).

**Fig 2:**
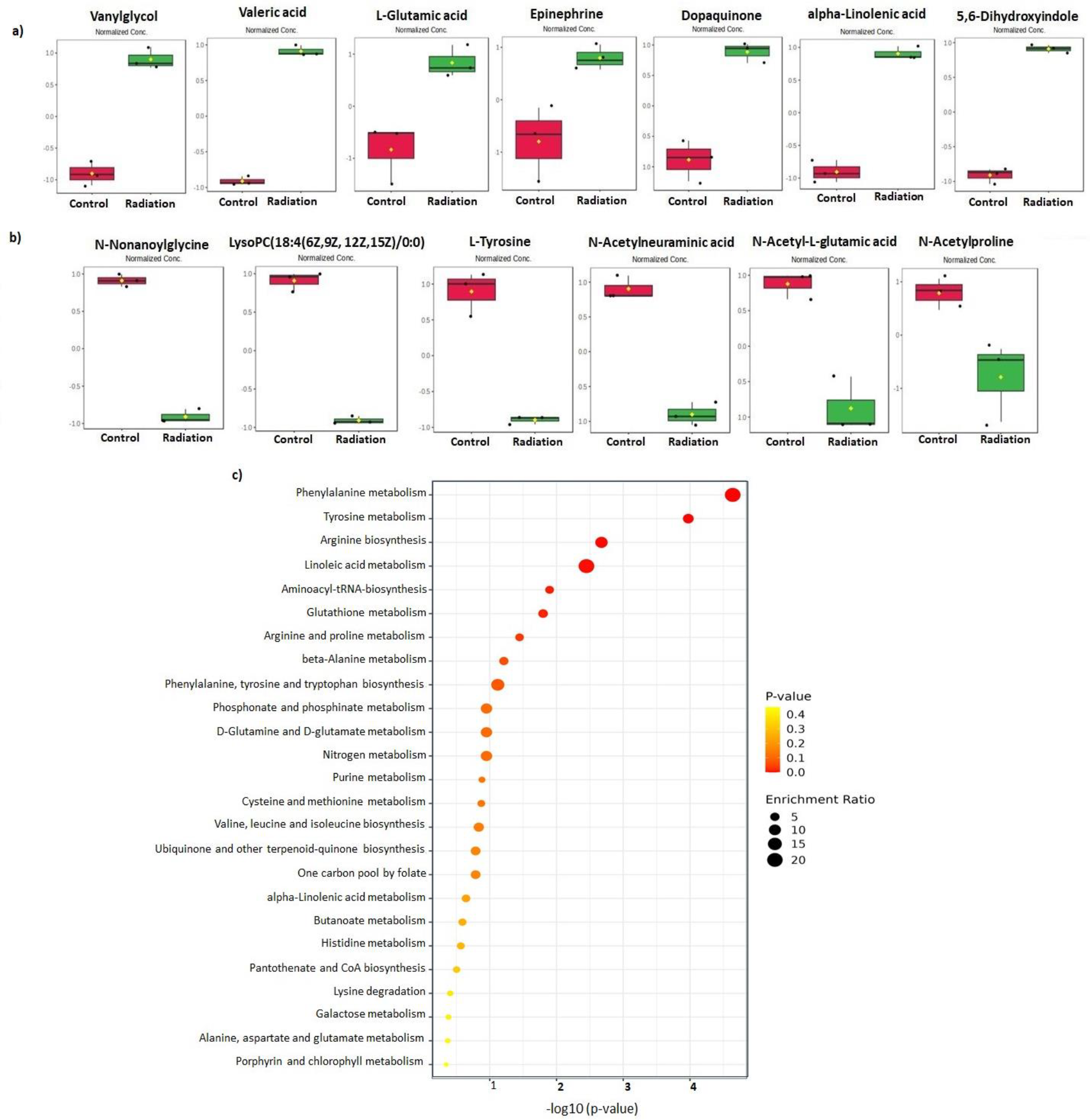
Effect of low-dose radiation exposure on hippocampal metabolites and pathways. a) Box plots of selected metabolites upregulated in radiation; b) Box-plots of selected metabolites downregulated in radiation; c) Different metabolomic pathways enriched by significantly altered metabolites post radiation exposure.

### Pathway enrichment analysis

To understand the influence of these metabolites on different pathways perturbed in neuronal diseases, we carried out pathway enrichment analysis based on the KEGG database. Phenylalanine metabolism, tyrosine metabolism, arginine biosynthesis, linoleic acid metabolism, aminoacyl t-RNA biosynthesis, and glutathione metabolism were among the top altered pathways (Fig 2c). Furthermore, we employed Metabolanalyst to perform joint pathway analysis where tyrosine metabolism was revealed as the most altered metabolism, along with ether lipid metabolism, pyrimidine metabolism, and steroid hormone biosynthesis emerging as the top altered pathways.

### Metabolomic and transcriptomic integrated analysis

To understand the association between transcriptomic regulation and metabolite levels, we performed multiomics analysis by integrating the list of significant metabolites obtained in the study with the set of differentially expressed genes (DEGs) set between the same two groups of animals obtained from a set of experimental hippocampal transcriptome study using the online tool Metabridge. Four genes in the input transcriptomic list were positively correlated with four metabolites. The alcohol dehydrogenase 7 (*Adh7*) gene encoding the alcohol dehydrogenase enzyme was associated with the metabolite 3,4-dihydroxyphenylglycol, which is involved in biological processes such as phosphatidylcholine biosynthesis, lipid transport, cell proliferation, and apoptosis. Glutaminase 2 (*Gls2*), which is involved in regulating glutaminase enzymes, is correlated with the metabolite glutamic acid, which is involved in excitatory neurotransmission and neuroinflammatory response (Ding et al. 2021). The SET and MYND domain containing 1 (*Smyd1*) gene, which regulates the enzyme [histone H3]-lysine4 N-trimethyltransferase, was correlated with the metabolite S-adenosylhomocysteine, involved in the production of homocysteine that plays a crucial role in maintaining neuronal homeostasis. Valyl-tRNA synthetase 1 (*Vars1*), which regulates the action of valine-tRNA ligase, was correlated with the metabolite L-valine. Further, integrated transcriptomic and metabolomic analysis using MetScape revealed that tyrosine metabolism was among the majorly affected pathways, along with other pathways such as purine and pyrimidine metabolism, valine, leucine, and isoleucine degradation, phenylalanine metabolism, alanine, aspartate, and glutamate metabolism.

We further performed a network analysis of the interaction of metabolites altered in the radiation group with the interacting metabolites and the same set of DEGs using Metscape. Among the different pathways altered, tyrosine metabolism and arginine, proline, glutamate, and aspartate metabolism involved the interaction of major resultant metabolites (Fig 3a and 3b). Metabolites from our list, such as 4-fumarylacetoacetate, 5,6-dihydroxyindole, dopaquinone, L-tyrosine, phenethylamine, phenylpyruvate, phenethylamine, L-noradrenaline, L-adrenaline, phenylacetaldehyde, phenylacetic acid, 4-hydroxyphenylacetate, 3,4-dihydroxyphenylethyleneglycol along with other interacting metabolites were involved in tyrosine metabolism. Additionally, the network also displayed the set of genes in the neighborhood that are also involved in their regulation and functioning, among which the genes aldehyde dehydrogenase 1 family member A3 (*Aldh1a3)*, dopamine beta-hydroxylase (*Dbh)*, inositol monophosphatase 2 *(Impa2)*, pantothenate kinase 1 (*Pank1)*, hyaluronan synthase 1 *(Has1)*, ectonucleoside triphosphate diphosphohydrolase 1 *(Entpd1)*, ubiquitin specific peptidase 44 *(Usp44)*, carbonyl reductase 4 *(Cbr4)*, klotho (*Kl)*, cytochrome P450, family 7, subfamily a, and polypeptide 1 *(Cyp7a1)* were also differentially expressed in our transcriptomic data. Similarly, metabolites such as L-arginine, L-citruline, N-acetyl-L-aspartate, L-glutamate, 5-oxoproline, N-acetyl-L-glutamate and spermidine from our list of metabolites obtained were observed to interact with the associated metabolites jointly regulating arginine, proline, glutamate, aspartate and arginine metabolism. Here, *Aldh1a3*, along with other interacting genes, was also observed to regulate the metabolites of this pathway.

**Fig 3:**
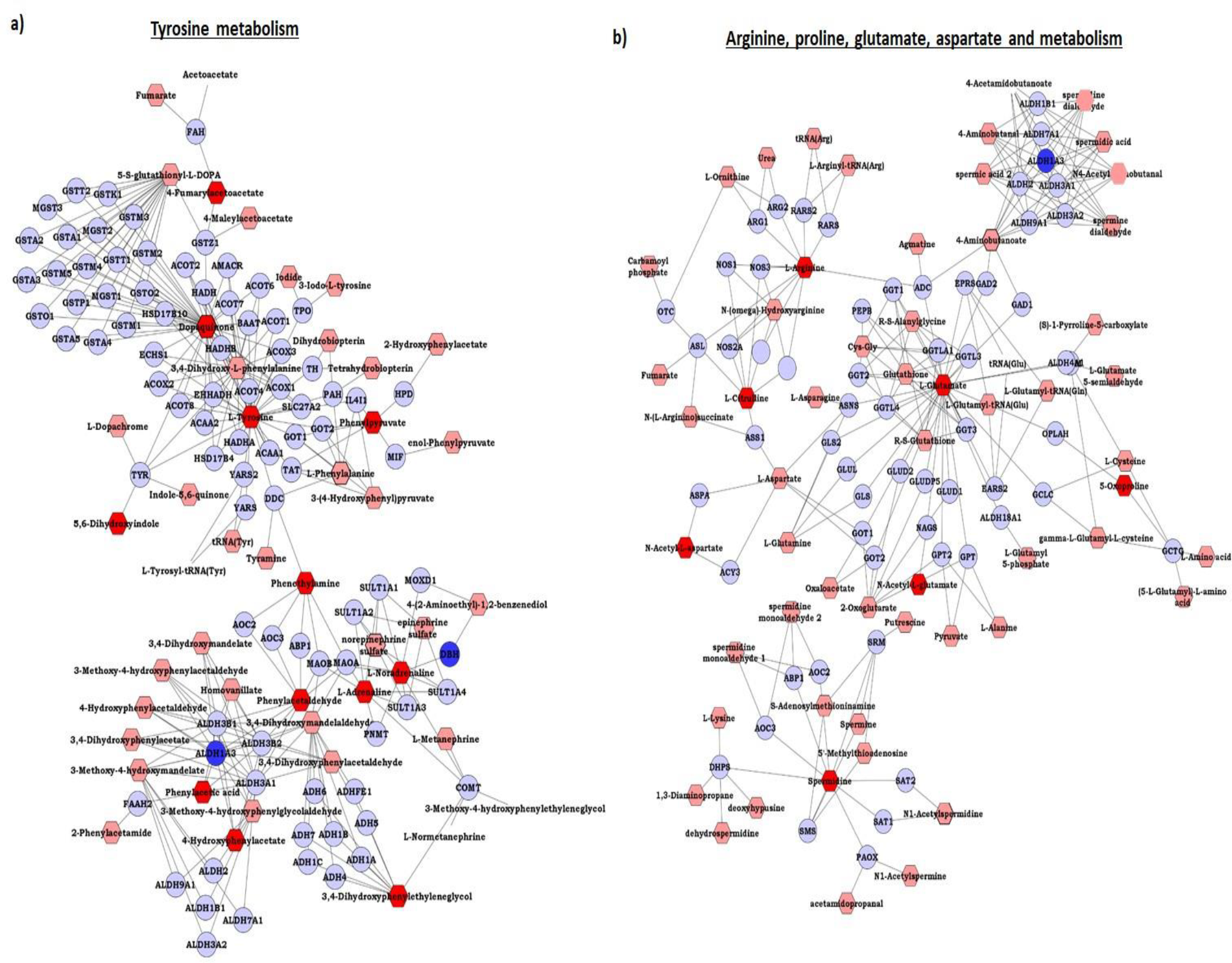
Effect of low-dose radiation exposure on amino acid metabolism. a) Network diagram of tyrosine metabolism and its interacting metabolites and genes b) Network diagram of arginine, proline, glutamate, and aspartate metabolism and its interacting metabolites and genes.

## Discussion

The present study employed untargeted metabolomics approach to scan the alterations of various metabolites in response to near-low dose radiation exposure in the hippocampus to understand its further implications and the pathways affected. One month old mice were exposed to 0.5 Gy X-ray, and the metabolomic profiling of the hippocampus was carried out to understand the acute changes in the brain metabolomic machinery. Our findings suggest that near low dose radiation can majorly alter amino acid levels and closely linked pathways such as tyrosine metabolism, arginine and proline metabolism, and glutamine metabolism, alluding to its involvement in neurotransmitter synthesis and the body’s antioxidant response.

The connectivity between genes and metabolites is an endless cyclic process that initiates with genes encoding a particular protein that, in turn, degrades into metabolites or makes use of metabolites for post-translational modifications and cell signaling processes that again assemble the same set of genes and proteins. Therefore, it is crucial to study the regulation of metabolite levels in response to various external factors due to their active participation in different cellular processes. Numerous studies have successfully employed metabolomics to garner detailed insights into the mechanisms prevailing in neurodegenerative disorders (Schumacher-Schuh et al. 2022). Recently, the use of a multiomics approach for comprehensive analysis of transcriptomic, proteomic, and metabolomic changes triggered post-exposure to xenobiotics has taken precedence.

Low-dose radiation exposure is often unavoidable in cases of diagnostic or therapeutic screening and administration. However, over the years, more individuals have reported undergoing such exposures, leading to concern among scientists regarding its long-term implications, particularly in neurodegenerative disorders (Rodgers et al. 2020). Additionally, such exposures in children are even more concerning as their brain is still developing and vulnerable to damage. IR exposure causes impairment in behavior and cognition, changes in neuronal morphology and death, and induces several alterations at the transcriptomic level (Narasimhamurthy et al. 2023). However, studies investigating metabolomic regulation after low-dose radiation exposure in the hippocampus are very limited. Radiation exposure resulted in the deterioration of the levels of L-tyrosine, N-acetyl-L-glutamic acid, N-acetylproline and N-nonanoylglycine, which are crucial amino acid metabolites that act as precursors in the synthesis of neurotransmitters. The impaired levels of these metabolites have been previously linked to neurodegeneration (Ayeni et al. 2022). On the other hand, amino acids and their derivatives, such as N-acetylglutamine, vanylglycol, valeric acid, N-undecanoylglycine, pyroglutamic acid, and L-glutamic acid, play an essential role in GABA neurotransmitter synthesis, fatty acid metabolism and glutathione metabolism. Epinephrine, a well-known neurotransmitter and hormone, was also elevated, while dopaquinone, a metabolite of L-DOPA known for facilitating the survival of dopaminergic neurons and crucial in cases of Parkinson’s like symptoms, was also elevated (Bruning et al. 2019). However, chronic elevated levels of dopaquinone are known to induce toxicity.

Pathway enrichment of the significantly affected metabolites revealed alterations in phenylalanine, tyrosine, arginine, and glutathione metabolism, among many others. These amino acid synthesis pathways are involved in various neurotransmitter and disease pathways. Tyrosine metabolism directly regulates the synthesis of L-DOPA to dopamine and indirectly regulates the synthesis of norepinephrine (Herman et al. 2019). Alterations in phenylalanine can cause modifications in the levels of catecholamine neurotransmitters (Fernstrom and Fernstrom 2007), while altered arginine metabolism has been linked to tauopathy (Mein et al. 2022). Furthermore, glutathione metabolism is crucial in providing defense against ROS and protecting neurons (Dringen et al. 2000). It should be noted that abnormal amino acid ratios and metabolism have been reported in the early stages of Alzheimer’s disease (Griffin and Bradshaw 2017) indicating that alterations in these metabolites can have serious neurodegenerative implications. To date, studies reporting altered amino acid metabolism in low-dose radiation are very sparse; however, Yamaguchi et al. have shown changes in a few amino acid residues and sequences in individuals exposed to low-dose radiation (Yamaguchi et al. 2022). Moreover, given the connection between low-dose radiation and neurotoxicity, as well as the demonstrated involvement of these metabolites and pathways in cases of neurotoxicity contributing to neurodegeneration, it can be inferred that low-dose radiation might trigger neurotoxicity by perturbing various amino acid metabolic pathways.

Multiomics integrated analysis helps find converging pathways and the crosstalk between different genes, enzymes, and metabolites. A previous study employing integrated metabolomics-DNA methylation analysis identified upregulated total amino acid synthesis, while downregulated aminoacyl-tRNA synthetase methylation was found (Torres et al. 2019). Another study involving joint transcriptomic and metabolomic analysis revealed nucleotide, amino acid, carbohydrate, lipid, and fatty acid metabolism, thus emphasizing the utility of multiomics approaches (Maan et al. 2023). Joint pathway analysis using Metaboanalyst revealed tyrosine metabolism, ether lipid metabolism, pyrimidine metabolism, and steroid hormone biosynthesis as the most jointly affected pathways. While the role of tyrosine metabolism is already emphasized, ether lipid metabolism is crucial for maintaining membrane integrity, taking part in various signaling pathways and differentiation, and is part of the antioxidant defense mechanism with a crucial role in various neurological diseases (Jové et al. 2023). Pyrimidine metabolism has been closely connected to mitochondrial function and is implicated in the pathogenesis of Alzheimer’s disease (Pesini et al. 2019). Furthermore, *de novo* pyrimidine synthesis in neural and glial cells is an essential process that, when affected, can affect signaling and neurotransmitter release and can lead to several neuropathies (Löffler et al. 2018). Hippocampal pyramidal neurons and glial cells have been demonstrated to be a site of neurosteroid hormone biosynthesis and have been involved in physiological functions such as neuronal growth and neuronal transmission, indicating the role of steroid hormones in the brain (Tsutsui et al. 2000).

The current study thus provides insight into the metabolomic response post-exposure to low doses of radiation, which is characterized by alterations in several amino acid synthesis pathways, glutathione metabolism, fatty acid biosynthesis, metabolism, etc., that play a crucial role in inducing neurotoxicity. Our study enhances the understanding of the link between their exposure and the manifestation of neurotoxicity in neurodegenerative-like conditions through a metabolomic approach and further employs integrated transcriptomic and metabolomic analysis to provide a holistic outlook and facilitate an enhanced understanding of the pathogenetic mechanism behind the damage associated with near low-dose exposure to radiation.

## Statements and Declarations

## Acknowledgments

The authors would like to thank the Director, Manipal School of Life Sciences, and Manipal Academy of Higher Education (MAHE) for support and infrastructure facilities. The authors would like to thank Dr. Gireesh G for his support throughout the study and his valuable suggestions for improving the manuscript and Mr. Sampara Vasishta for his help during metabolomic data analysis. RKN would like to thank MAHE, Manipal, for the Dr. T.M.A Pai fellowship and KSTePS, Govt of Karnataka, for the DST-PhD scholarship. This work was supported by The Science and Engineering Research Board [Grant No: ECR/2017/001239/LS]

## Funding

This work was supported by The Science and Engineering Research Board [Grant No: ECR/2017/001239/LS].

### Author Contributions

KDM conceptualized the study, KDM, BSSR designed the study, RKN maintained the animals, performed and analyzed the experiment, and wrote the manuscript. KS planned and helped perform the irradiation procedure. MBJ planned and helped with the mass spectrometry sample run. VBS, MBJ, KDM, BSSR and KS provided significant comments and revisions to the article.

### Ethics approval

Animals were obtained from the Central Animal Research Facility, Kasturba Medical College, Manipal Academy of Higher Education, Manipal, after duly obtaining ethical clearance from the Institutional Animal Ethics Committee (IAEC/KMC/108/2019), Manipal Academy of Higher Education, Manipal.

### Declaration of Competing Interests

The authors declare that the research was conducted in the absence of any commercial or financial relationships that could be construed as a potential conflict of interest.

## Notes

### Competing Interest Statement

The authors have declared no competing interest.

