## Supplementary material for "Deciphering the Metabolic Shifts in The Hippocampus of Mice Subjected to Near Low Dose Radiation: Insights from Metabolomics and Integrated Multi-omics": table 1

**Table 1: List of significantly altered metabolites in the radiation group with respect to the control with their fold change values**

| Metabolite | Fold Change | log2(FC) | p.value | FDR |
|---|---|---|---|---|
| Valeric acid | 0.0054409 | -7.5219 | 0.00000515 | 0.00045 |
| 5,6-Dihydroxyindole | 0.019874 | -5.653 | 1.30E-05 | 0.00048 |
| (S)-3-Octanol glucoside | 0.02087 | -5.5824 | 0.00030152 | 0.00325 |
| 2-Oxo-4-methylthiobutanoic acid | 0.025179 | -5.3116 | 0.00022837 | 0.00309 |
| 2-Hydroxy-3-methylpentanoic acid | 0.026428 | -5.2418 | 0.0004263 | 0.00332 |
| alpha-Linolenic acid | 0.030362 | -5.0416 | 8.91E-05 | 0.00196 |
| Adenine | 0.035241 | -4.8266 | 0.0030415 | 0.00937 |
| Phosphorylcholine | 0.039706 | -4.6545 | 0.00039696 | 0.00332 |
| p-Cresol | 0.042352 | -4.5614 | 0.000132 | 0.00235 |
| Dictyoquinazol C | 0.043434 | -4.525 | 0.0012021 | 0.00578 |
| 3-Methoxytyrosine | 0.049623 | -4.3328 | 0.00288 | 0.00937 |
| o-Cresol | 0.058073 | -4.106 | 0.00029744 | 0.00325 |
| Phenylacetic acid | 0.063031 | -3.9878 | 0.0016699 | 0.007 |
| Vanylglycol | 0.06417 | -3.9619 | 0.00028421 | 0.00325 |
| Adipic acid | 0.068305 | -3.8719 | 0.0029413 | 0.00937 |
| Aminomalonic acid | 0.081447 | -3.618 | 0.00327 | 0.00937 |
| p-Hydroxyphenylacetic acid | 0.085801 | -3.5429 | 0.0020205 | 0.0079 |
| Ethyl 3-mercaptobutyrate | 0.088168 | -3.5036 | 0.0039957 | 0.01082 |
| 3,4-Methylenesebacic acid | 0.10253 | -3.2858 | 0.0033 | 0.00937 |
| Quinone | 0.10634 | -3.2333 | 0.00054995 | 0.00346 |
| 13-HODE | 0.13008 | -2.9425 | 0.03134 | 0.04955 |
| 9'-Carboxy-gamma-chromanol | 0.13961 | -2.8405 | 0.0013449 | 0.00621 |
| m-Cresol | 0.1426 | -2.81 | 0.0023984 | 0.00844 |
| Succinyladenosine | 0.14841 | -2.7523 | 0.005636 | 0.01329 |
| CPA(18:2(9Z,12Z)/0:0) | 0.15036 | -2.7335 | 0.021698 | 0.03637 |
| Phenytoin catechol | 0.16418 | -2.6066 | 0.002978 | 0.00937 |
| MG(20:5(5Z,8Z,11Z,14Z,17Z)/0:0/0:0) | 0.16653 | -2.5862 | 0.0032492 | 0.00937 |
| MG(0:0/20:5(5Z,8Z,11Z,14Z,17Z)/0:0) | 0.16738 | -2.5788 | 0.0031976 | 0.00937 |
| alpha-Linolenic acid | 0.16756 | -2.5772 | 0.00036778 | 0.00332 |
| Dopaquinone | 0.16899 | -2.565 | 0.001215 | 0.00578 |
| Oxypurinol | 0.17997 | -2.4742 | 0.0014895 | 0.00655 |
| Epinephrine | 0.18652 | -2.4226 | 0.022861 | 0.03796 |
| 12-Ketoporrigenin | 0.19659 | -2.3467 | 0.00013353 | 0.00235 |
| Indoleacrylic acid | 0.20277 | -2.3021 | 0.027444 | 0.04431 |
| 3-Phenylpropyl isovalerate | 0.23472 | -2.091 | 0.0087793 | 0.01753 |
| Ethyl 4-phenylbutanoate | 0.23496 | -2.0895 | 0.0029811 | 0.00937 |
| 5,10-Methylene-THF | 0.24123 | -2.0515 | 0.0088649 | 0.01753 |

Draft

| | | | | |
|---|---|---|---|---|
| Phenyllactic acid | 0.27971 | -1.838 | 0.0056652 | 0.01329 |
| 7-Methylguanine | 0.30645 | -1.7063 | 0.019135 | 0.0327 |
| 2-Oxo-4-methylthiobutanoic acid | 0.30779 | -1.7 | 0.001578 | 0.00677 |
| p-Cresol glucuronide | 0.34531 | -1.534 | 0.0036845 | 0.01013 |
| Oleamide | 0.35222 | -1.5054 | 0.01407 | 0.02553 |
| Malic acid | 0.38336 | -1.3832 | 0.013035 | 0.02415 |
| Phenylethylamine | 0.38678 | -1.3704 | 0.0087045 | 0.01753 |
| Phenylacetaldehyde | 0.42163 | -1.246 | 0.029099 | 0.04656 |
| L-Glutamic acid | 0.42561 | -1.2324 | 0.01091 | 0.02043 |
| Glutarylcarnitine | 2.1294 | 1.0905 | 0.032164 | 0.04955 |
| N6,N6,N6-Trimethyl-L-lysine | 2.1969 | 1.1355 | 0.0030034 | 0.00937 |
| 2-Pyrroloylglycine | 2.4237 | 1.2772 | 0.003222 | 0.00937 |
| Monomenthyl succinate | 2.4527 | 1.2943 | 0.001148 | 0.00577 |
| Neuroprotectin D1 | 2.5176 | 1.3321 | 0.0066495 | 0.01481 |
| L-Arginine | 2.6055 | 1.3816 | 0.032374 | 0.04955 |
| 2-Keto-6-acetamidocaproate | 2.6319 | 1.3961 | 0.010633 | 0.02012 |
| 9,10-Epoxyoctadecenoic acid | 2.7003 | 1.4331 | 0.016485 | 0.02873 |
| N-Desmethyldiphenhydramine | 3.0381 | 1.6032 | 0.0081103 | 0.0166 |
| Spermidine | 3.1159 | 1.6396 | 0.032159 | 0.04955 |
| L-Leucine | 3.4968 | 1.806 | 0.00043329 | 0.00332 |
| dinor-Levomethadyl acetate | 3.554 | 1.8295 | 0.0068001 | 0.01496 |
| 5-Methyldeoxycytidine | 3.5915 | 1.8446 | 0.00053495 | 0.00346 |
| Methyl 3-(methylthio)butanoate | 3.7294 | 1.8989 | 0.0077743 | 0.01628 |
| Adenosine | 4.0796 | 2.0284 | 0.010541 | 0.02012 |
| 3-Oxododecanoic acid | 4.4347 | 2.1488 | 7.66E-07 | 0.00013 |
| Octadecyl fumarate | 4.4664 | 2.1591 | 0.031577 | 0.04955 |
| 3,4-Dihydroxyphenylglycol | 4.99 | 2.3191 | 0.013288 | 0.02436 |
| 2-Hexenoylcarnitine | 5.0358 | 2.3322 | 0.0046478 | 0.01186 |
| N-Acetylneuraminic acid | 5.0923 | 2.3483 | 0.00019179 | 0.00281 |
| 14,15-DiHETrE | 6.0364 | 2.5937 | 0.0070047 | 0.01522 |
| N-Acetylproline | 6.2287 | 2.6389 | 0.02658 | 0.04332 |
| L-Tyrosine | 6.2677 | 2.6479 | 0.00057529 | 0.00349 |
| N-Nonanoylglycine | 6.2887 | 2.6528 | 1.37E-05 | 0.00048 |
| Galactitol | 6.514 | 2.7035 | 0.0004848 | 0.00341 |
| 17-HDoHE | 6.5995 | 2.7224 | 0.00041653 | 0.00332 |
| PE-NMe(13D5/13D5) | 6.6461 | 2.7325 | 0.00017132 | 0.00274 |
| 13-L-Hydroperoxylinoleic acid | 6.6736 | 2.7385 | 0.0078263 | 0.01628 |
| Creatinine | 6.9874 | 2.8048 | 0.0063581 | 0.01447 |
| Methyl sorbate | 7.1886 | 2.8457 | 0.0013767 | 0.00621 |
| PS(DiMe(13,5)/MonoMe(13,5)) | 7.3347 | 2.8747 | 0.00059435 | 0.00349 |

Draft

| | | | | |
|---|---|---|---|---|
| L-Citramalyl-CoA | 7.4205 | 2.8915 | 0.0010164 | 0.00526 |
| Vanillactic acid | 7.8014 | 2.9637 | 0.020759 | 0.03513 |
| Hydroxyvalerylcarnitine | 8.9704 | 3.1652 | 0.0041357 | 0.01103 |
| Phenylacetylglutamine | 9.0142 | 3.1722 | 0.0064109 | 0.01447 |
| Pyroglutamylglycine | 9.0847 | 3.1834 | 0.016381 | 0.02873 |
| Isobutyryl-L-carnitine | 9.585 | 3.2608 | 0.0017833 | 0.00713 |
| Guanosine | 9.6673 | 3.2731 | 0.00069188 | 0.00393 |
| Dodecanedioylcarnitine | 9.6704 | 3.2736 | 0.0077623 | 0.01628 |
| Resolvin D5 | 9.6888 | 3.2763 | 0.00054369 | 0.00346 |
| PE(14:1(9Z)/14:0) | 10.621 | 3.4089 | 0.0022556 | 0.00827 |
| 13-L-Hydroperoxylinoleic acid | 10.911 | 3.4478 | 0.00039538 | 0.00332 |
| PA(18:1(11Z)/14:0) | 12.104 | 3.5975 | 0.0044019 | 0.01139 |
| 19(20)-EpDPE | 12.524 | 3.6466 | 9.14E-06 | 0.00048 |
| PG(i-12:0/18:2(9Z,11Z)) | 12.755 | 3.673 | 0.0047362 | 0.01191 |
| 3-hydroxydecanoyl carnitine | 12.785 | 3.6764 | 0.0020827 | 0.00797 |
| PA(16:0/14:0) | 12.845 | 3.6831 | 0.0050334 | 0.01248 |
| PA(8:0/22:0) | 13.051 | 3.7061 | 0.0052033 | 0.01272 |
| 4,8 Dimethylnonanoyl carnitine | 13.057 | 3.7068 | 8.81E-05 | 0.00196 |
| Prostaglandin J2 | 13.32 | 3.7355 | 0.0078631 | 0.01628 |
| Pyroglutamic acid | 13.56 | 3.7613 | 0.0017622 | 0.00713 |
| N-Undecanoylglycine | 19.292 | 4.2699 | 0.0063597 | 0.01447 |
| 5Z-Dodecenoic acid | 19.295 | 4.2701 | 0.010439 | 0.02012 |
| N-Acetyl-L-glutamic acid | 26.734 | 4.7406 | 0.0021647 | 0.00811 |
| N-Acetylglutamine | 56.425 | 5.8183 | 0.0028583 | 0.00937 |
| LysoPA(24:1(15Z)/0:0) | 80.227 | 6.326 | 0.00083946 | 0.00461 |
| LysoPC(18:4(6Z,9Z,12Z,15Z)/0:0) | 81.411 | 6.3471 | 1.98E-05 | 0.00058 |

Draft
